# Deep learning-guided design of cell type-specific AAV promoters

**DOI:** 10.64898/2026.01.13.699371

**Authors:** Sean K. Wang, Boxiong Deng, Surag Nair, Xiaobai Ren, Jiaying Li, Jenna Tijerina, Praveen Prakhar, Ziming Luo, Chelsea Nnebe, Samuel H. Kim, Yimin Zhou, Sahil H. Shah, Alexander Davis, Rohan Mahajan, Yilin Qiao, Yuxi Zhou, Jenny Zhang, Yunlu Xue, Jeffrey L. Goldberg, Wei Wei, Anshul Kundaje, Howard Y. Chang, Sui Wang

**Author notes:** These authors contributed equally to this work (Sean K. Wang, Boxiong Deng).

## Abstract

Precise cell type targeting is critical for both clinical and experimental applications of adeno-associated viral (AAV) vectors, yet engineering vectors with cell type-specific activity remains a challenge. Here, we compared three strategies leveraging single-cell chromatin accessibility data to design cell type-specific AAV promoters, including a deep learning-based method to generate *de novo* regulatory sequences. When applied to target retinal ganglion cells or horizontal cells in mouse retina, deep learning-guided design consistently outperformed rational approaches, yielding synthetic promoters with stronger and more specific expression *in vivo*. Synthetic AAV promoters supported diverse transgenes, enabling the recording and ablation of targeted cells. Promoter activity was also maintained in human retinal organoids, suggesting that deep learning-designed sequences may be suitable for translation. Our findings highlight the potential of deep learning to synthesize cell type-specific AAV promoters and establish a versatile platform for cell type targeting with broad implications for gene therapy and basic research.

## INTRODUCTION

Adeno-associated viral (AAV) vectors are central to many emerging strategies to treat genetic diseases, including gene replacement therapy, gene editing, gene silencing, and the expression of protective genes^1–5^. Beyond their clinical applications, AAV vectors are also powerful research tools, enabling detailed investigations of gene function, mapping of neural circuitry, and the exploration of fundamental biological processes^6,7^. Precise targeting of AAV vectors to the intended cell types is essential for achieving therapeutic efficacy and ensuring accurate mechanistic insights. Conversely, off-target expression of AAV vectors may lead to cellular toxicity and immune responses that compromise both experimental validity and patient safety^8–10^.

Despite the importance of cell type specificity, targeted AAV vectors are still lacking for many cell types. One potential solution is to use cell type-specific AAV promoters that confine expression to the desired cells. However, designing such promoters remains a significant challenge. A common design strategy is to clone the region upstream of a cell type-specific gene with the aim of recapitulating its endogenous expression, but native regulatory sequences often span multiple kilobases (kb) and must be substantially shortened to meet the size constraints of AAV vectors^11–13^. Alternative methods to generate promoters include combining conserved sequence elements, transcription factor binding sites (TFBSs), or open chromatin regions, yet these options frequently fail to attain cell type specificity^14–16^. For example, in a screen of 230 rationally designed AAV promoters targeting retinal cell types, only 17 (7.4%) exhibited cell type specificity better than 90%^14^, highlighting the need for a different approach.

Deep learning has emerged as a powerful tool for decoding the *cis*-regulatory logic of the genome by identifying the intricate sequence patterns that underlie gene expression^17–19^. At the same time, deep learning-based methods are rapidly advancing synthetic biology, enabling the design of *de novo* sequences ranging from novel regulatory elements to fully functional proteins^20–24^. Previously, we trained deep learning models derived from the BPNet architecture on single-cell assay for transposase-accessible chromatin sequencing (scATAC-seq) data from human retinas to predict the regulatory activity of variants implicated in eye diseases^18,25^. Similar models of chromatin accessibility recently succeeded in creating cell type-specific enhancers in *Drosophila* by constructing sequences with predicted activity in only the targeted cells^26,27^. We thus wondered if deep learning could also be applied to obtain cell type-specific AAV promoters.

In this study, we compared three different strategies to design cell type-specific AAV promoters from single-cell chromatin accessibility data, including a deep learning-based method to generate *de novo* regulatory sequences. To evaluate these strategies, we targeted two cell types in the mammalian retina: retinal ganglion cells (RGCs) and horizontal cells (HCs). AAV promoters designed by deep learning consistently outperformed those generated by rational approaches, achieving stronger and more specific expression in vivo. These findings demonstrate the potential of deep learning to synthesize AAV promoters targeting specific cell types, offering a versatile platform for both gene therapy and basic research.

## RESULTS

### Design of AAV promoters using cell type-specific regulatory sequences

We initially attempted to create an AAV promoter targeting HCs in the retina by employing the 2 kb region upstream of mouse *Lhx1*, a HC-specific gene^28^. However, this promoter failed to achieve HC specificity in mice (Supplementary Fig. 1a,b), highlighting the limitations of using native regulatory sequences for cell type targeting. To design cell type-specific AAV promoters, we instead leveraged cell type chromatin accessibility profiles from scATAC-seq and devised three strategies based on these data (Fig. 1a). In the first strategy, we incorporated genomic regions with high accessibility in only the target cell type, reasoning that the selective accessibility of these elements might confer cell type-specific expression. In the second strategy, we incorporated scATAC-seq peaks enriched in cell type-specific TFBSs, an approach we previously showed can identify active *cis*-regulatory regions^29^. In the third strategy, we incorporated *de novo* sequences predicted by deep learning models of chromatin accessibility to be uniquely accessible in the target cell type. We termed the resulting sequences from these three strategies “candidate enhancers” and combined them with the base promoter of a cell type-specific gene to form full-length AAV promoters.

**Figure 1.**
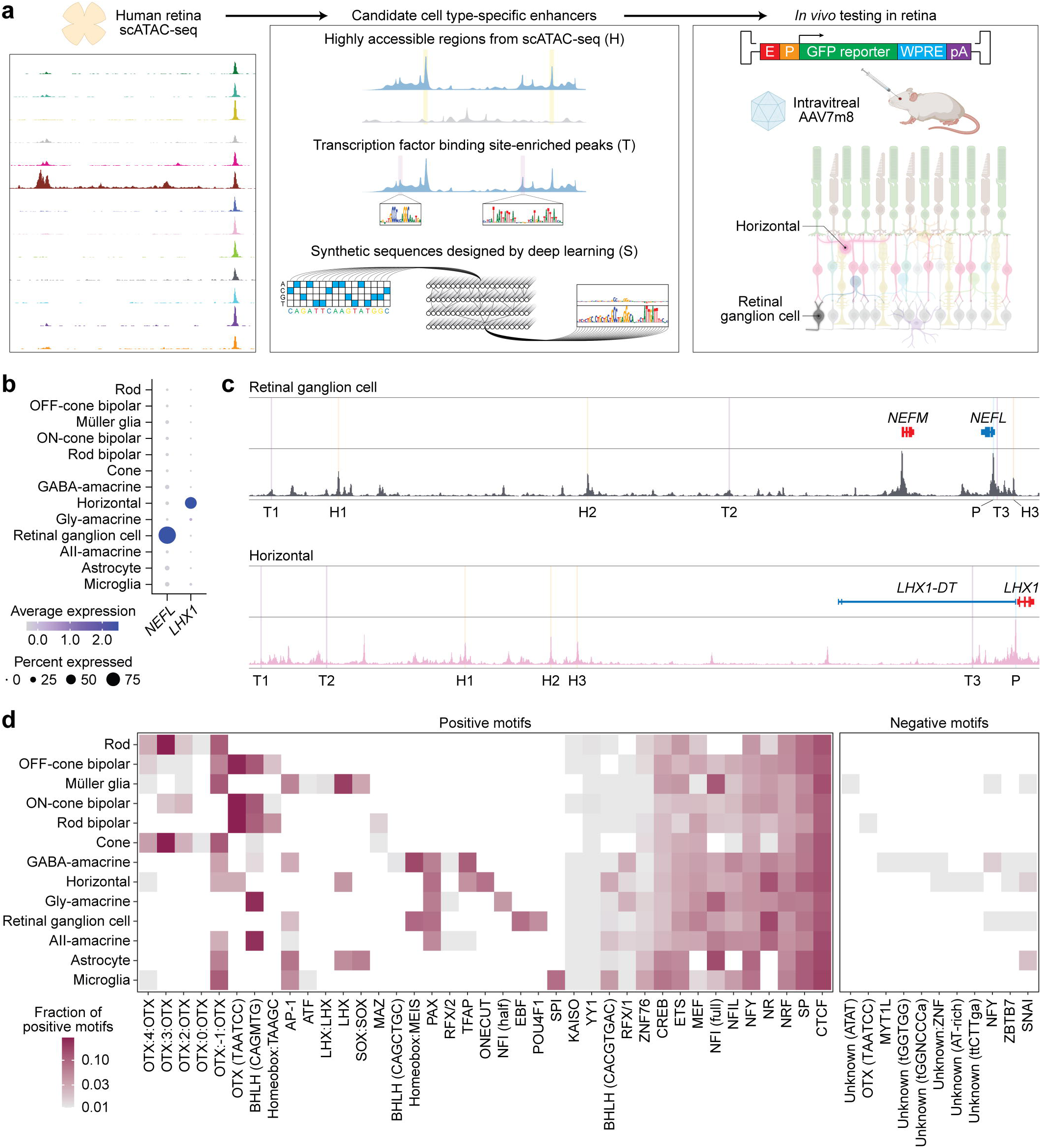
AAV promoter design strategies. a, Overview of AAV promoter design strategies. Candidate cell type-specific enhancers were designed using human retina scATAC-seq data and consisted of 1) highly accessible regions (H), 2) TFBS-enriched peaks (T), and 3) synthetic sequences designed by deep learning (S). AAV promoters composed of a candidate enhancer (E) joined to the base promoter (P) of a cell type-specific gene were placed upstream of a GFP reporter sequence, packaged in the AAV7m8 capsid, and intravitreally injected into mice to target RGCs or HCs in the retina. **b,** Dot plot visualizing the normalized RNA expression of selected genes in human retinal cell types^25^. The color and size of each dot correspond to the average expression level and fraction of expressing cells, respectively. **c,** Sequencing tracks of chromatin accessibility in RGCs or HCs. Each track depicts the aggregate scATAC-seq signal in the given cell type normalized by the number of reads in TSS regions. Genes transcribed in the sense direction (TSS on the left) are shown in red, and genes in the antisense direction (TSS on the right) are shown in blue. Vertical lines indicate locations of the base promoter (P), highly accessible regions (H1-H3), and TFBS-enriched peaks (T1-T3) selected to target each cell type. Coordinates for each region: retinal ganglion cell (chr8:24607917-24978286), horizontal (chr17:36612867-36946800). **d,** Heatmap of sequence motifs positively or negatively associated with chromatin accessibility in human retinal cell types. Motifs were identified from deep learning models using TF-MoDISco^49^. Color corresponds to the frequency of each motif as a fraction of all positive motif instances in that cell type. For OTX:n:OTX motifs, n refers to the number of intervening bases.

We applied the above strategies to generate AAV promoters targeting RGCs and HCs. We first identified a base promoter for each cell type by comparing the expression and accessibility of putative marker genes using single-cell RNA sequencing (scRNA-seq) and scATAC-seq data from human retina^25^. *NEFL* and *LHX1* exhibited cell type-specific expression and accessibility in RGCs and HCs, respectively (Fig 1b. and Supplementary Fig. 2a-d), leading us to select a 500 bp region upstream of *NEFL* and 411 bp region upstream of *LHX1* as base promoters. Using human retina scATAC-seq data, we next located three highly accessible regions (H1-H3) and three TFBS-enriched peaks (T1-T3) in each cell type (Fig. 1c and Supplementary Fig. 3a,b). These endogenous candidate enhancers were 500 bp in length, yielding total promoter sizes of 1,000 bp for RGCs and 911 bp for HCs.

To generate *de novo* sequences targeting RGCs and HCs, we used BPNet deep learning models trained on pseudo-bulk scATAC-seq data from 13 human retinal cell types^18,25^. These models can predict the cell type accessibility of DNA sequences in the retina and were tasked with creating candidate enhancers uniquely accessible in RGCs or HCs. We began by extracting sequence motifs from BPNet models and determining those positively or negatively associated with chromatin accessibility in each cell type (Fig. 1d). Starting with a 170 bp sequence for RGCs or 320 bp sequence for HCs, we then iteratively added or removed motifs and introduced single-base mutations to maximize the difference in predicted accessibility between target and off-target cells. We repeated this process to design 100 candidate sequences per target cell type and scored their enhancer activity *in silico* across random genomic regions. The three top scoring sequences were selected as synthetic enhancers (S1-S3), and combining these enhancers with base promoters yielded total promoter sizes of 670 bp for RGCs and 731 bp for HCs.

### Targeting of retinal ganglion cells

AAV promoters were placed upstream of a green fluorescent protein (GFP) reporter gene and packaged in the AAV7m8 capsid, which efficiently transduces the retina following intravitreal delivery in mice (Supplementary Fig. 4a-c)^30^. We first assessed AAV promoters targeting RGCs, the primary cell type affected in glaucoma and other optic neuropathies^31^. Testing of the *NEFL* base promoter (RGC_P) in mouse retina demonstrated minimal GFP expression in RBPMS-positive RGCs (Fig. 2a), indicating that this sequence alone was insufficient to drive robust target cell activity. Of the three promoters incorporating regions highly accessible in RGCs, two showed GFP expression in RGCs, but also in off-target amacrine cells (ACs) (Fig. 2b and Supplementary Fig. 5a). Similarly, all three promoters incorporating scATAC-seq peaks enriched in RGC-specific TFBSs displayed activity in both RGCs and ACs (Fig. 2b and Supplementary Fig. 5b). In contrast, all three AAV promoters incorporating synthetic sequences designed by deep learning drove GFP expression specifically in RGCs (Fig. 2b and Supplementary Fig. 5c), suggesting this strategy to be the best for achieving cell type specificity.

**Figure 2.**
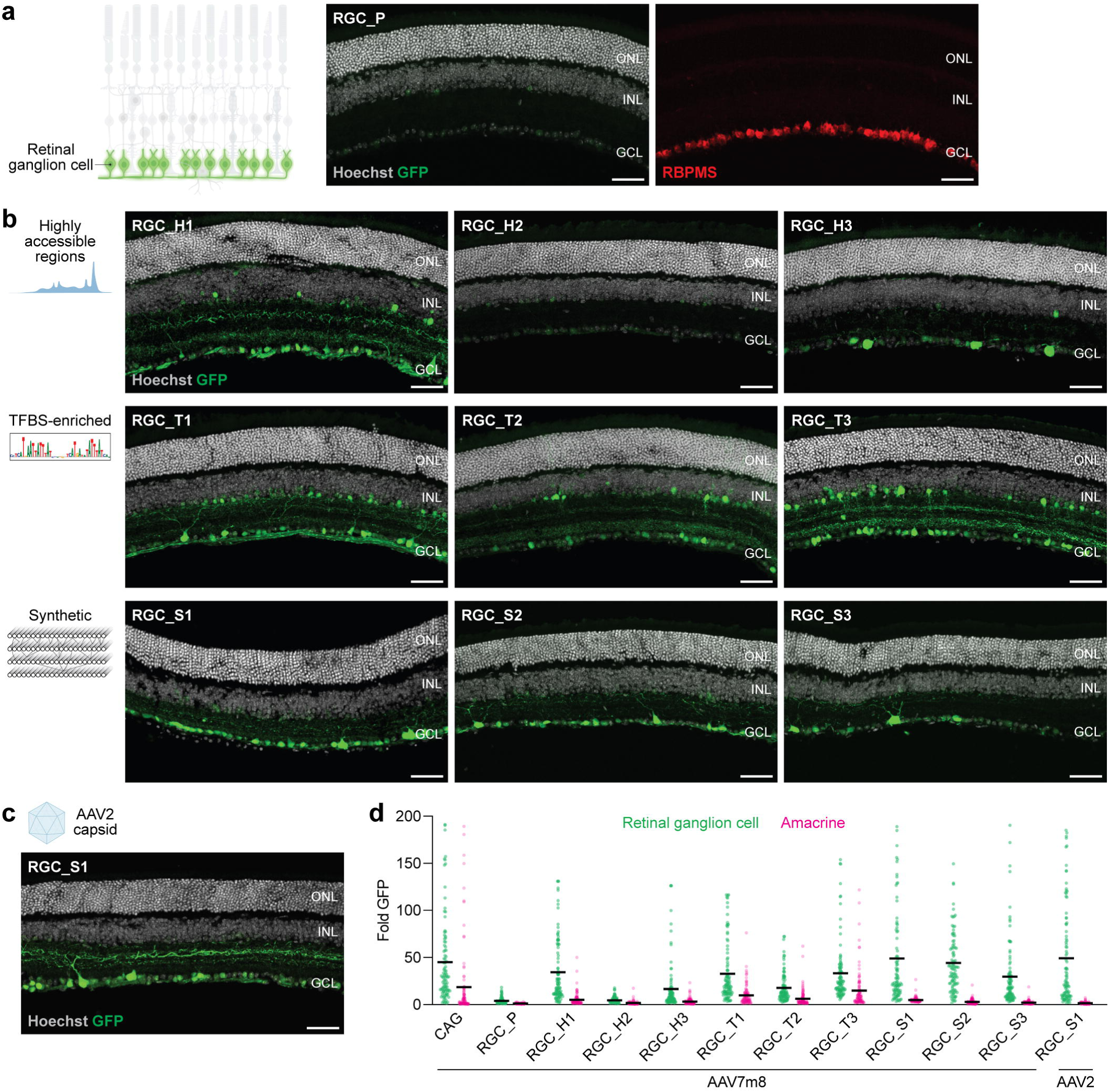
Synthetic AAV promoters enable retinal ganglion cell-specific expression. a, *Left:* Schematic of RGCs in the retina. *Right:* RGC_P activity and RBPMS immunostaining in mouse retina three weeks after AAV7m8 transduction. n = 3 eyes. Scale bars, 50 μm. ONL, outer nuclear layer; INL, inner nuclear layer; GCL, ganglion cell layer. **b,** Activity of the indicated AAV promoters in mouse retina three weeks after AAV7m8 transduction. n = 3-6 eyes per group. Scale bars, 50 μm. **c,** RGC_S1 promoter activity in mouse retina three weeks after AAV2 transduction. n = 3 eyes. Scale bar, 50 μm. **d,** Quantification of AAV promoter activity in RGCs and ACs. Each point represents a cell, and black lines depict means. Values for GFP expression are presented relative to that in the outer nuclear layer and are not shown if greater than 200-fold. n = 100 of each cell type from 3-6 eyes per group.

Because RGCs reside in the inner retina, we additionally evaluated the activity of the RGC_S1 synthetic promoter when packaged in the AAV2 capsid (Fig. 2c and Supplementary Fig. 5d), which exhibits more restrictive tropism than AAV7m8 following intravitreal delivery^30^. Quantification of GFP fluorescence revealed this capsid and promoter combination to drive strong and specific expression in RGCs (Fig 2d).

### Targeting of horizontal cells

We next compared AAV promoters targeting HCs, a group of interneurons that mediate lateral inhibition in retinal circuits^32^. Prior screening of rationally designed AAV promoters in the retina failed to identify a specific promoter for this cell type^14^, underscoring the difficulty of attaining HC-specific expression. Initial testing of the *LHX1* base promoter (HC_P) in mice showed no detectable activity in CALB1-positive HCs (Fig. 3a). For all three promoters incorporating regions highly accessible in HCs, GFP expression was seen in off-target RGCs and ACs, but not in HCs (Fig. 3b and Supplementary Fig. 6a). This was likewise the case for two of the three promoters incorporating scATAC-seq peaks enriched in HC-specific TFBSs, with only HC_T1 yielding rare GFP-positive HCs (Fig. 3b and Supplementary Fig. 6b). Conversely, all three AAV promoters incorporating synthetic sequences designed by deep learning demonstrated HC expression (Fig. 3b and Supplementary Fig. 6c), again suggesting this strategy to be the most effective for cell type targeting.

**Figure 3.**
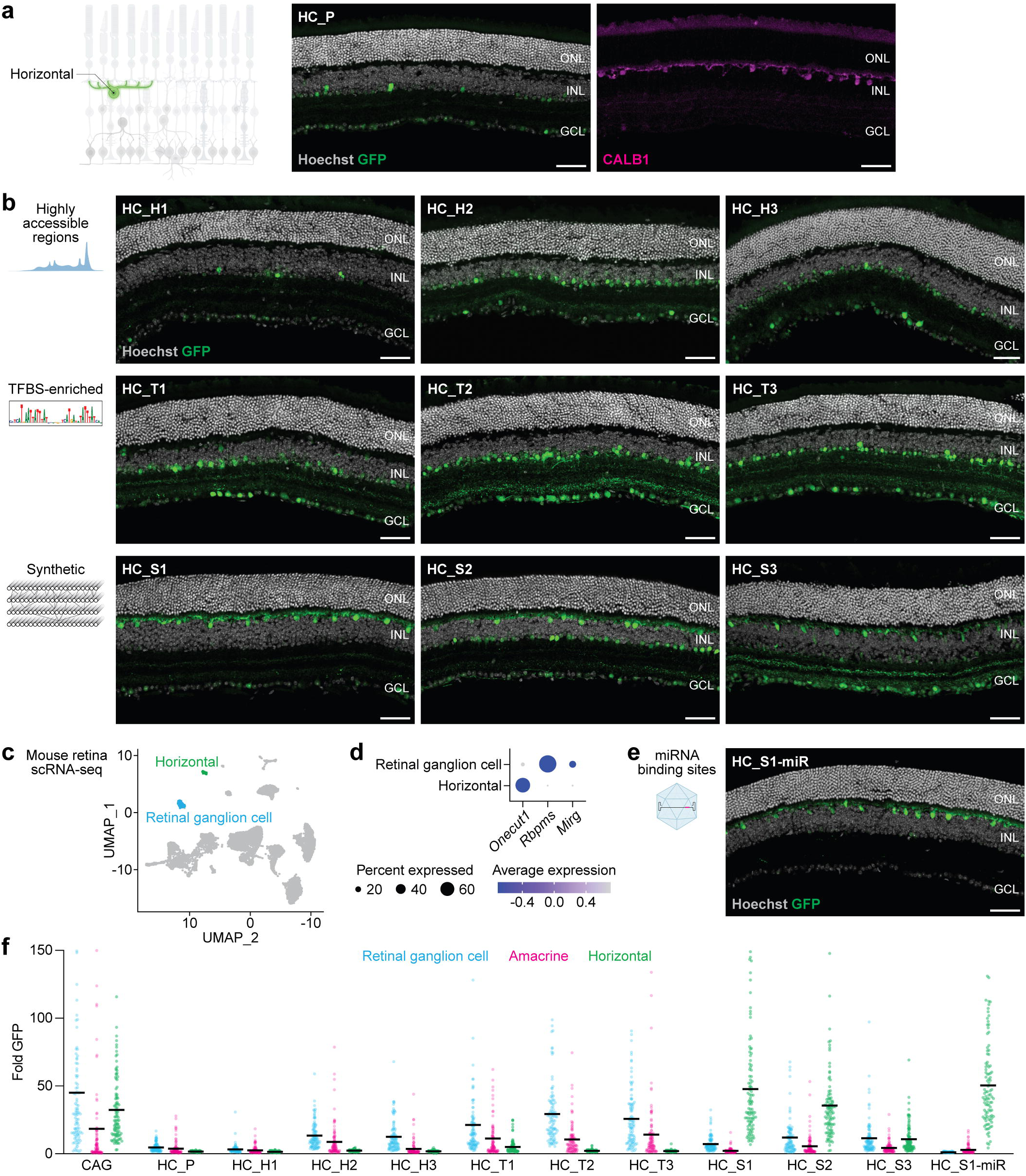
Synthetic AAV promoters enable horizontal cell-specific expression. a, *Left:* Schematic of HCs in the retina. *Right:* HC_P activity and CALB1 immunostaining in mouse retina three weeks after AAV7m8 transduction. n = 3 eyes. Scale bars, 50 μm. ONL, outer nuclear layer; INL, inner nuclear layer; GCL, ganglion cell layer. **b,** Activity of the indicated AAV promoters in mouse retina three weeks after AAV7m8 transduction. n = 3-6 eyes per group. Scale bars, 50 μm. **c,** Uniform manifold approximation and projection (UMAP) of mouse retina cells from scRNA-seq performed by Macosko et al^33^. **d,** Dot plot visualizing the normalized RNA expression of selected genes in RGCs and HCs. **e,** HC_S1-miR promoter activity in mouse retina three weeks after AAV7m8 transduction. n = 6 eyes. Scale bar, 50 μm. **f,** Quantification of AAV promoter activity in RGCs, ACs, and HCs. Each point represents a cell, and black lines depict means. Values for GFP expression are presented relative to that in the outer nuclear layer and are not shown if greater than 150-fold. n = 100 of each cell type from 3-6 eyes per group.

The HC_S1 synthetic promoter exhibited the strongest activity in HCs, but also displayed weak off-target expression in RGCs. We hypothesized that this off-target activity could be mitigated by leveraging endogenous microRNAs to silence RGC expression. Using mouse scRNA-seq data^33^, we searched for microRNAs expressed in RGCs but not HCs and identified one such cluster of microRNAs located within a gene called *Mirg* (Fig. 3c,d). We selected five microRNAs (miR-134-5p, miR-154-5p, miR-299a-5p, miR-323-3p, and miR-379-5p) within *Mirg* expressed in neural tissues and fully conserved between mouse and human and introduced binding sites for them in the vector genome^34^. Addition of microRNA binding sites to HC_S1 (HC_S1-miR) eliminated GFP fluorescence from RGCs, resulting in HC-specific expression (Fig. 3e,f and Supplementary Fig. 6d). For challenging cell types, the specificity of synthetic AAV promoters can thus be refined by using microRNAs to suppress off-target activity.

### Functional applications of synthetic AAV promoters

Genetically encoded calcium indicators such as jGCaMP8m enable recording of neuronal activity by visualizing changes in intracellular calcium^35^. Using the RGC_S1 promoter, we generated an AAV vector expressing jGCaMP8m in RGCs and recorded the responses of individual cells to visual stimuli (Fig. 4a,b). Following intravitreal delivery of the vector in mice, explanted retinas were presented with a bright bar moving in different directions and calcium responses imaged using two-photon microscopy. Transduced cells exhibited a wide range of response profiles consistent with the functional diversity of RGCs^36^, including sustained or transient On and Off responses (Fig. 4c) as well as direction- and orientation-selective activity (Fig. 4d). These results show that RGC_S1 drives expression in RGCs spanning diverse functional classes.

**Figure 4.**
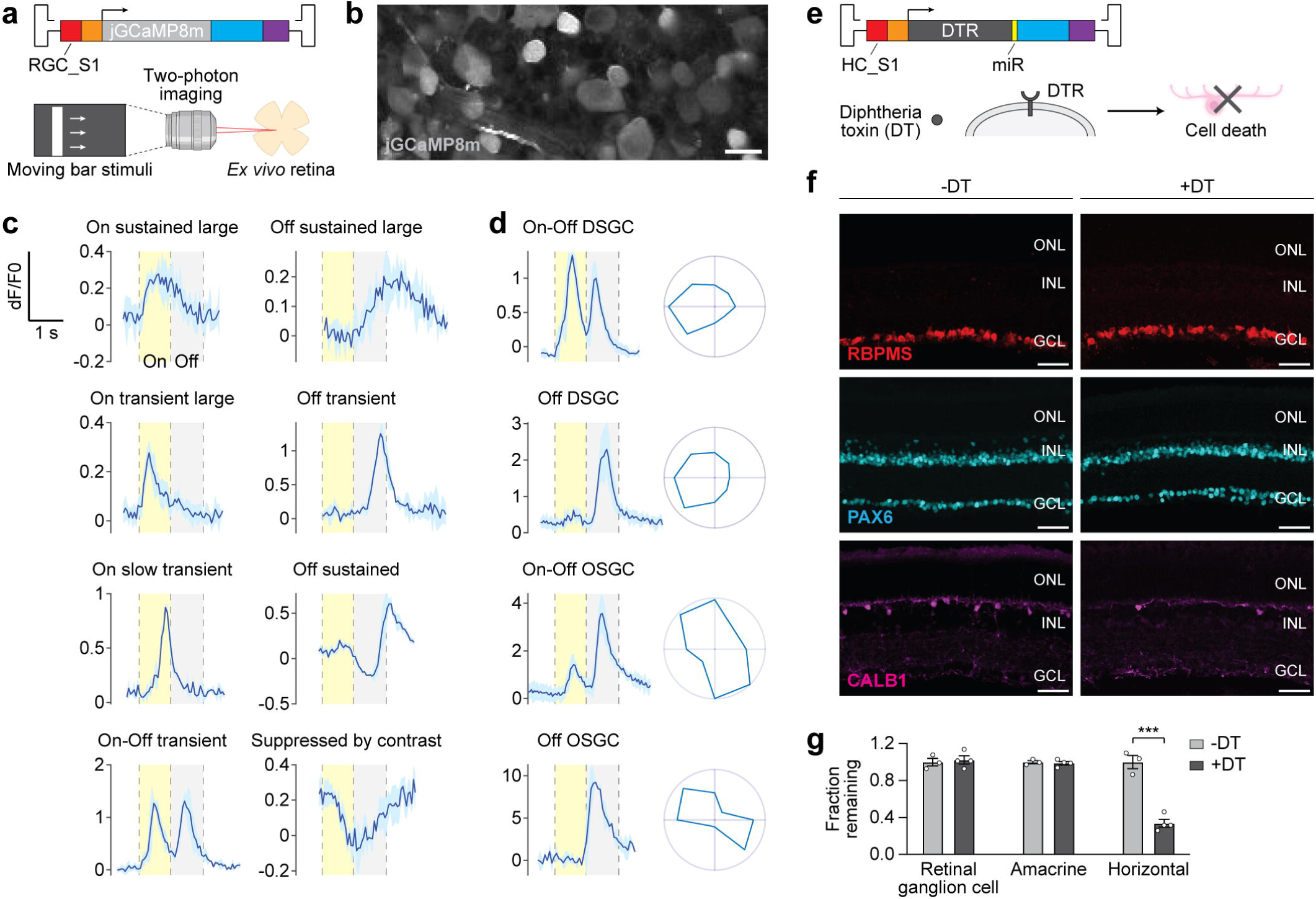
Synthetic AAV promoters enable cell type-specific recording and ablation. a, Schematic of calcium imaging experiments to record RGC activity. Moving bar stimuli were projected onto *ex vivo* mouse retinas three weeks after intravitreal injection of AAV7m8-RGC_S1-jGCaMP8m, and jGCaMP8m responses measured in RGCs using two-photon imaging. **b,** jGCaMP8m expression in the ganglion cell layer. n = 6 eyes. Scale bar, 20 μm. **c,** Example jGCaMP8m responses to moving bar stimuli in different RGC subtypes. Dashed lines separate On and Off response windows. Traces represent mean (dark blue) ± SD (light blue). n = 3-5 recordings per cell from 6 eyes. **d,** *Left:* Example jGCaMP8m responses in direction-selective ganglion cells (DSGCs) and orientation-selective ganglion cells (OSGCs). *Right:* Polar plots of peak responses for cells shown on left. Traces represent mean (dark blue) ± SD (light blue). n = 3-5 recordings per cell from 6 eyes. **e,** Schematic of diphtheria toxin (DT) experiments to ablate HCs. Mice were intravitreally injected with AAV7m8-HC_S1-DTR-miR encoding diphtheria toxin receptor (DTR), then intravitreally injected with DT after two weeks. **f,** RBPMS, PAX6, and CALB1 immunostaining in mouse retina three weeks after AAV transduction with or without diphtheria toxin. n = 3-4 eyes per group. Scale bars, 50 μm. ONL, outer nuclear layer; INL, inner nuclear layer; GCL, ganglion cell layer. **g,** Quantification of RGC, AC, and HC ablation. Data are mean ± SEM. n = 3-4 eyes per group. *** *P*<0.001 by two-tailed Student’s t test.

We also tested if synthetic promoter expression of diphtheria toxin receptor (DTR) could mediate targeted cell ablation in the retina. In the presence of diphtheria toxin, cells expressing DTR undergo irreversible inhibition of protein synthesis, resulting in cell death^37^. Conversely, wild-type mouse cells are resistant to diphtheria toxin even at high doses. Using HC_S1-miR, we generated an AAV vector expressing DTR in mouse HCs and compared transduced eyes with and without diphtheria toxin (Fig. 4e). Following a single intravitreal injection of diphtheria toxin, we observed a 66.1% decline in HCs (*P*<0.001) but no significant change in the number of RGCs or ACs (Fig. 4f,g), indicating HC-specific ablation. Overall, these data highlight the versatility of synthetic AAV promoters as tools for cell type-specific interrogation and manipulation.

### Synthetic promoter activity in human retinal organoids

We lastly assessed the translational relevance of synthetic AAV promoters by testing their activity in human retinal organoids. Retinal organoids were derived from Brn3b-tdTomato human embryonic stem cells (hESCs) (Fig. 5a), which express the Brn3b marker of RGCs fused with tdTomato^38^. This modification enabled identification of differentiated RGCs by tdTomato expression. To identify HCs, we performed immunostaining for the marker protein PROX1^39^.

**Figure 5.**
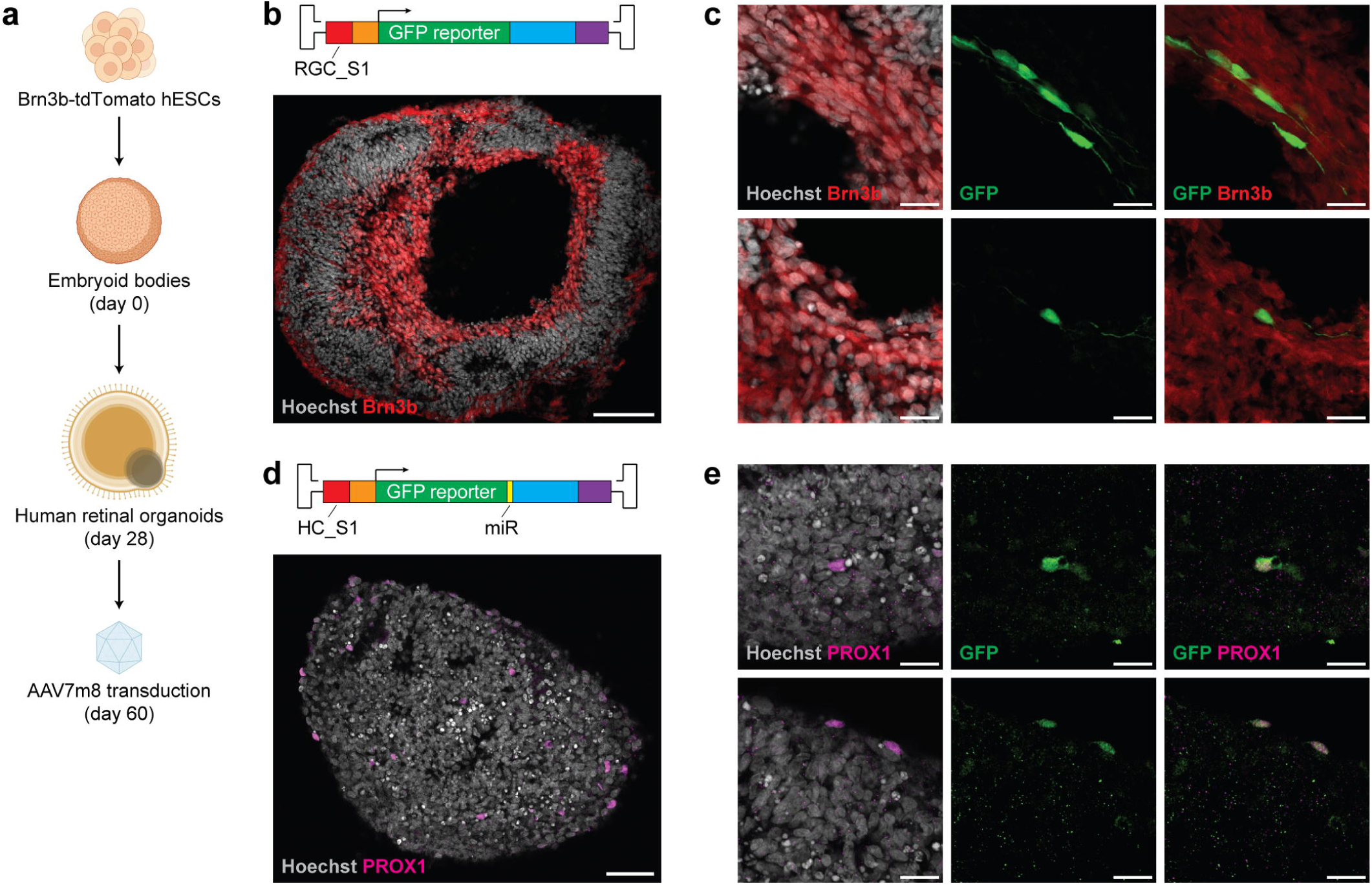
Synthetic AAV promoters drive expression in human retinal organoids. a, Schematic of human retinal organoid experiments. Human embryonic stem cells (hESCs) expressing tdTomato under control of the Brn3b (*POU4F2*) locus were induced to form embryoid bodies followed by differentiation into human retinal organoids. AAV7m8 transduction was performed at day 60 of differentiation. **b,** Brn3b-tdTomato expression in human retinal organoid. n = 3 organoids. Scale bar, 100 μm. **c,** RGC_S1 promoter activity and Brn3b-tdTomato expression in human retinal organoid one week after AAV transduction. n = 3 organoids. Scale bars, 20 μm. **d,** PROX1 immunostaining in human retinal organoid. n = 3 organoids. Scale bar, 50 μm. **e,** HC_S1-miR promoter activity and PROX1 immunostaining in human retinal organoid one week after AAV transduction. n = 3 organoids. Scale bars, 20 μm.

Retinal organoids were treated with AAV vectors at day 60 of differentiation, around the peak of RGC abundance^40^. Organoids at this time point displayed tdTomato fluorescence predominantly in the inner retina, consistent with the distribution expected for RGCs (Fig. 5b). Although AAV transduction efficiency in organoids was low, we detected RGC_S1-driven GFP expression in tdTomato-positive RGCs (Fig. 5c). Similarly, HC_S1-miR-driven GFP expression in retinal organoids was seen in PROX1-positive HCs (Fig. 5d,e). Together, our findings suggest that synthetic AAV promoters designed using human chromatin accessibility data may retain cell type-specific activity in humans.

## DISCUSSION

Precise cell type targeting is critical for both clinical and experimental applications of AAV vectors, yet achieving robust cell type-specific expression remains challenging. Here, we compared three strategies to engineer cell type-specific AAV promoters and found that synthetic sequences designed by deep learning consistently outperformed rationally selected regulatory elements *in vivo*. Using this method, we generated AAV promoters driving strong and specific expression in RGCs, a therapeutically relevant cell type, as well as HCs, a cell type for which targeted promoters are lacking. Synthetic AAV promoters supported the expression of diverse transgenes, such as jGCaMP8m for cell type-specific recording and DTR for cell type-specific ablation. Importantly, the activity of these promoters was maintained in human retinal organoids, suggesting that AAV promoters designed by models trained on human accessibility data may be suitable for translation. In principle, our deep learning approach can be applied to the vast array of cell types for which chromatin accessibility data are available.

*De novo* enhancers generated by deep learning were 170 or 320 bp in length, yielding synthetic promoter sizes of 670 bp for RGC_S1 and 731 bp for HC-S1 (839 bp including microRNA binding sites). Although these sizes are well within the range of commonly used AAV promoters, it is possible that they could be shortened to accommodate larger transgenes. Taskiran et al. recently described synthetic enhancers as small as 49 bp that conferred cell type specificity in *Drosophila*^26^, and Yin et al. reported regulatory elements of comparable length in human cell lines^22^. In line with these results, model interpretation of our synthetic enhancer sequences revealed stretches of bases with minimal contribution to predicted accessibility, suggesting that these regions could potentially be trimmed while preserving cell type-specific activity. Notably, for certain motifs such as OTX, the spacing between motif instances was associated with differential accessibility across cell types. Deep learning models in our study thus captured not only the vocabulary of sequence motifs driving cell type accessibility, but also the grammar of their interactions.

Our work complements advancements in AAV capsid engineering that have greatly enhanced the tropism of vectors for specific tissues, but are insufficient on their own to restrict activity to individual cell types^30,41,42^. Combining modified capsids with deep learning-guided promoter design may enable more precise targeting than either of these approaches alone, reducing risks from off-target expression and permitting lower doses to be used. *De novo* enhancers from our method might also facilitate control of large gene cargoes through transcriptional crosstalk of AAV concatemers^43^. This phenomenon allows regulatory regions on one AAV genome to direct expression from another, expanding the utility of the cell type-specific sequences that we can design. Our study additionally builds upon emerging deep learning-based strategies to create synthetic regulatory elements from gene expression data^20,22,44^. These methods may provide a parallel route to optimize the strength and specificity of AAV promoters and could potentially be integrated with accessibility-based models to improve performance. Finally, generation of synthetic AAV promoters will benefit from ongoing efforts to construct comprehensive single-cell chromatin accessibility atlases of human tissues^45,46^, which offer a rich source of training data for regulatory sequence design. By extending the framework described here to these datasets, it may be possible to engineer vectors targeting the full spectrum of human cell types.

## METHODS

### AAV promoter design

AAV promoters were designed by combining candidate enhancers with the base promoter of a cell type-specific gene. For each target cell type, a base promoter was first selected by comparing the expression of ten putative marker genes using human retina scRNA-seq data and identifying the three most specific genes detected in at least 50% of target cells^25^. From these genes, the promoter region whose accessibility was most specific to the target cell type was chosen. MACS2 was then used to call peaks within this region from the target cell fragments file^25,47^, and the nearest peak to the TSS was designated as the base promoter. For HCs, a 411 bp peak 1.2 kb upstream of the *LHX1* TSS was designated as the base promoter. For RGCs, the nearest peak to the *NEFL* TSS overlapped the gene’s coding sequence; therefore, the 500 bp region immediately upstream of the *NEFL* start codon was designated as the base promoter.

Candidate cell type-specific enhancers were designed using three different strategies leveraging human retina scATAC-seq data. Endogenous candidate enhancers were selected from scATAC-seq marker peaks located within 1 megabase (Mb) of the base promoter. For each peak, the log2 fold change in accessibility compared to non-target cell types was divided by the false discovery rate, and the three peaks with the largest result were designated as highly accessible regions (H1-H3). For each marker peak, the number of cell type-specific TFBSs was also calculated using the searchSeq function in TFBSTools^48^. Cell type-specific TFBSs were identified from previous motif enrichment analysis of scATAC-seq peaks and included EBF1, ESRRA, ESRRB, ESRRG, NR1D1, NR1D2, NR2F2, NR4A1, NR4A3, NR5A1, NR5A2, POU1F1, POU4F1, POU4F3, RORA, RORB, and RORC for RGCs and KLF5, KLF16, ONECUT1, TFAP2A, TFAP2B, TFAP2C, ZNF148, and ZNF263 for HCs^25^. Non-overlapping TFBSs with scores >10 were counted, and the three peaks with the greatest number of TFBSs were designated as TFBS-enriched peaks (T1-T3).

Synthetic cell type-specific enhancers were designed using previously described BPNet models (https://zenodo.org/records/6330053) that predict chromatin accessibility for human retinal cell types from DNA sequence^18,25^. These models take a 2,114 bp sequence as input and predict the accessibility profile (as a probability distribution) and natural log of the total ATAC-seq counts (as a scalar value) in the central 1,000 bp. For each cell type, five model folds were trained on different chromosome splits. Each fold was used to predict accessibility counts on 10,000 cell type peaks and 10,000 GC-matched non-peak regions to obtain a distribution of predicted counts over accessible and inaccessible sequences. Accessibility counts were then normalized from 0 to 1 using the empirical cumulative distribution function.

Candidate sequences were generated using a motif-based optimization strategy with the objective defined as the normalized accessibility in the target cell type minus the highest normalized accessibility among non-target cell types. Sequence motifs from fold 0 models were first extracted using TF-MoDISco and clustered across cell types using gimmemotifs cluster^49,50^, resulting in a map of motifs positively or negatively associated with accessibility in each cell type. A list of curated motifs for each target cell type was then created by filtering for positive motifs found in the target and up to two non-target cell types, as well as negative motifs found only in non-target cell types. Sequences were designed by starting with a random 2,114 bp genomic region devoid of accessibility peaks and modifying the central 170 bp (for RGCs) or 320 bp (for HCs) while keeping flanking regions constant. Five rounds of optimization were performed, with each round consisting of one motif-insertion step and two single-base-mutation steps. During each motif-insertion step, consensus versions of curated motifs were inserted at 10 random positions in both forward and reverse orientations. During each single-base-mutation step, 10 random positions were mutated to each possible base. Following every modification, the sequence was scored using model fold 0, 1, or 2 chosen at random, and a beam search strategy was used to track the top three scoring sequences. At the end of five rounds of optimization, the central 170 or 320 bp of the top scoring sequence was added to a list of candidate sequences. This process was repeated 100 times to generate 100 candidate sequences for each target cell type.

Candidate sequences were ranked using held-out model folds 3 and 4. Each of the 100 sequences was embedded into 100 different 2,114 bp genomic regions inaccessible in the target cell type. Sequences were then ranked by the average normalized accessibility in the target cell type minus the highest average normalized accessibility among non-target cell types, and the three top scoring sequences were designated as synthetic enhancers (S1-S3).

### Plasmid construction

Base promoter and endogenous enhancer sequences were synthesized as gene fragments (Integrated DNA Technologies) or amplified from human genomic DNA using nested PCR. Synthetic enhancer sequences were synthesized as gene fragments or by annealing complementary oligonucleotides (Integrated DNA Technologies). Enhancer and base promoter sequences were cloned downstream of a 5’ inverted terminal repeat (ITR) and upstream of an EGFP-P2A-Cre coding sequence, woodchuck hepatitis virus posttranscriptional regulatory element (WPRE), bovine growth hormone polyadenylation signal, and 3’ ITR using Gibson assembly. For jGCaMP8m and DTR plasmids, EGFP-P2A-Cre was replaced with the coding sequence from pGP-CMV-jGCaMP8m (Addgene, #162372) or pAAV-FLEX-DTR-GFP (Addgene, #124364), respectively^35,51^.

### AAV vector production and delivery

Recombinant AAV8, AAV7m8, and AAV2 vectors were produced and titered as previously described^29^. Intravitreal injections were performed in 5-6-week-old CD-1 (Charles River Laboratories, #022) or C57BL/6 (The Jackson Laboratory, #000664) wild-type mice using a pulled glass needle connected to a Picospritzer microinjector (Parker Hannifin). Approximately 1 μL of AAV7m8 or AAV2 vector (10^12^-10^13^ genome copies per mL) was administered per eye. Subretinal injections were performed in postnatal day 0 C57BL/6 mice using a pulled glass needle connected to a FemtoJet microinjector (Eppendorf). Approximately 0.25 μL of AAV8 vector (10^12^-10^13^ genome copies per mL) was administered per eye. Both male and female mice were used and were randomly assigned to treatment groups unless indicated otherwise. All animal studies were approved by the Institutional Animal Care and Use Committee at Stanford University (APLAC-33948), Lingang Laboratory (NZXSP-2022-4), or the University of Chicago (ACUP-72247).

### Histology and immunohistochemistry

Mice were perfused as previously described^52^. Following perfusion, eyes were enucleated and fixed in 4% paraformaldehyde in phosphate-buffered saline (PBS) (pH 7.4) at room temperature for 1 hour. Retinas were then isolated in cold PBS for histology and immunostaining. For sections, retinas were equilibrated in a series of sucrose solutions, frozen in a 1:1 mixture of optimal cutting temperature compound (Tissue-Tek) and 30% sucrose in PBS, and cut on a cryostat (Leica Microsystems) at a thickness of 20 μm. Sections were washed in PBS, permeabilized in 0.2% Triton X-100 in PBS for 20 minutes, incubated in a blocking solution (0.1% Triton X-100, 1% bovine serum albumin, and 10% donkey serum in PBS) for 30 minutes, and stained with primary antibodies in blocking solution at room temperature for 2 hours or 4°C overnight. Sections were subsequently washed in 0.1% Triton X-100 in PBS, stained with secondary antibodies at room temperature for 30 minutes to 2 hours, and mounted onto glass slides with Fluoromount-G (Southern Biotechnology). For flat-mounts, whole retinas were immersed in a blocking solution (3% Triton X-100, 0.5% Tween-20, 1% bovine serum albumin, 0.1% sodium azide, and 10% donkey serum in PBS) at 4°C for 24 hours. Retinas were stained with primary antibodies in blocking solution at room temperature for 8 hours followed by 4°C for 48 hours. Retinas were subsequently washed in 0.1% Triton X-100 in PBS, stained with secondary antibodies at room temperature for 2 hours followed by 4°C for 24 hours, relaxed with four radial cuts, and mounted onto glass slides with the ganglion cell layer facing up.

Primary antibodies included guinea pig anti-RBPMS (PhosphoSolutions, 1832-RBPMS, 1:500), rabbit anti-PAX6 (Thermo Fisher Scientific, 42-6600, 1:500), mouse anti-CALB1 (Sigma-Aldrich, C9848, 1:500), and rabbit anti-PROX1 (BioLegend, 925202, 1:500). Secondary antibodies were obtained from Jackson ImmunoResearch and included Cy3 donkey anti-guinea pig (706-165-148), Alexa Fluor 647 donkey anti-rabbit (711-605-152), Cy3 donkey anti-rabbit (711-166-152), Alexa Fluor 647 donkey anti-mouse (715-601-151), and Alexa Fluor 594 donkey anti-mouse (715-585-151). Nuclei were labeled with 4′,6-diamidino-2-phenylindole (DAPI) (Sigma-Aldrich) or Hoechst 33342 (Thermo Fisher Scientific).

### Confocal imaging and analysis

Retinal images were acquired on an Olympus FV3000 confocal microscope using a 4x, 20x, or 40x air objective. For sections, z-stacks spanning 10-20 μm of tissue were captured. Maximum intensity projections of approximately 5 μm were used for representative images. For flat-mounts, z-stacks spanning 60-90 μm of tissue were captured. Maximum intensity projections of the ganglion cell, AC, and HC layers were used for representative images and identified by immunostaining for RBPMS, PAX6, and CALB1, respectively.

Image analysis was performed using Fiji (ImageJ). Regions of interest (ROIs) in retinal sections corresponding to background, individual cell soma, and the outer nuclear layer were manually selected. AAV promoter activity was quantified by subtracting fluorescence in the background region from raw fluorescence signals in ROIs, then dividing the mean GFP intensity in individual cell soma by that in the outer nuclear layer. RGCs, ACs, and HCs were defined as RBPMS-positive, PAX6-positive, and CALB1-positive cells, respectively.

### MicroRNA selection

Mouse retina scRNA-seq data from Macosko et al. was analyzed using Seurat^33,53^. After filtering for cells with >900 detected genes, gene expression counts were normalized using the NormalizeData function and scaled using the ScaleData function. Graph-based clustering was then performed using the top 20 principal components at a resolution of 0.5, and RGC and HC clusters annotated by their expression of cell type marker genes. Transcripts differentially expressed between RGCs and HCs were identified using the FindMarkers function.

MicroRNAs within *Mirg* were filtered for those expressed in neural tissues (adult mouse brain and induced neurons), but not adult kidney, heart, lung, or liver^34^. The remaining microRNAs were filtered for those fully conserved between mouse and human as determined by miRBase^54^, resulting in five retained sequences. Binding sites for the five microRNAs corresponding to their reverse complement sequences were tiled without intervening bases and cloned into the AAV plasmid between the transgene and WPRE.

### GCaMP-mediated optical recording

Mice injected with AAV7m8-RGC_S1-jGCaMP8m were euthanized after dark adaptation and retinas isolated in oxygenated Ames’ medium (Sigma-Aldrich) under infrared illumination. Retinas were maintained in darkness in room temperature Ames’ medium bubbled with 95% O_2_/5% CO_2_ for 0-3 hours until use. A white organic light-emitting display (OLED) (eMagin, 800 by 600 pixels, 60 Hz refresh rate) was presented to each retina at a resolution of 1.1 μm/pixel. Moving bar stimuli were generated using MATLAB and the Psychophysics Toolbox^55^ and projected through the condenser lens of a custom two-photon microscope (Bruker Nano Surfaces Division) onto the photoreceptor layer. A bright bar (220 µm wide, 440 µm long) with an intensity of ∼6.3×10^4^ isomerizations (R*)/rod/s and background intensity of ∼1.8x10^3^ R*/rod/s was moved along the long axis at a speed of 440 µm/s over a 660 µm diameter field. jGCaMP8m in RGCs was excited by a Chameleon Ultra II Ti:sapphire laser (Coherent) tuned to 920 nm, and visual stimuli were presented after 10 s of continuous laser scanning. To separate visual stimuli from jGCaMP8m fluorescence, a band-pass filter (Semrock) was placed on the OLED to pass light peaked at 470 nm, while two notched filters (Bruker Nano Surfaces Division) were placed before the photomultiplier tubes to block light of the same wavelength. jGCaMP8m responses were recorded at 13-20 Hz using a 60x water immersion objective (Olympus).

Recordings were analyzed using Fiji and MATLAB as previously described^56^. ROIs corresponding to background and soma were manually selected. Fluorescence in the background region was subtracted from raw fluorescence signals in ROIs, and the mean jGCaMP8m intensity in each ROI was determined for every frame. Measurements obtained between visual stimuli were used to create a baseline (F0) trace for each ROI. Fluorescence values (F) were converted to dF/F0 by calculating dF = (F-F0)/F0. dF/F0 traces were then clipped, sorted by visual stimulus direction (inward or outward), and averaged over trials. Peak dF/F0 values were calculated using custom MATLAB scripts.

### Diphtheria toxin-mediated cell ablation

Mice were intravitreally injected with 2 μL of diphtheria toxin (5 ng/μL) (Sigma-Aldrich) two weeks after AAV7m8-HC_S1-DTR-miR transduction to ablate cells expressing DTR. RGCs and HCs in retinal sections were quantified by counting the number of RBPMS-positive and CALB1-positive cells, respectively, in a single z-slice per eye. ACs in retinal sections were quantified by counting the number of PAX6-positive cells within the central 200 μm of a single z-slice per eye.

### Human retinal organoids

Brn3b-tdTomato hESCs (a gift from Dr. Donald Zack) were maintained under feeder-free conditions in StemFlex medium (Thermo Fisher Scientific) at 37 °C with 5% CO₂ and differentiated into retinal organoids as previously described^57^. Day 0 of differentiation was defined by the formation of embryoid bodies, and organoids were cultured in suspension beginning day 28. Day 60 retinal organoids were transduced with 10^10^ genome copies of AAV vector in a 50 µL volume, followed by 50/50 media exchanges every 3 days without further addition of vector. After 1 week, organoids were fixed in 4% paraformaldehyde at room temperature for 30 minutes, equilibrated and frozen as described above, and cut on a cryostat (Leica Microsystems) at a thickness of 20 μm.

### Sequencing tracks

Sequencing tracks of scATAC-seq data from Wang et al. were visualized using the WashU Epigenome Browser^25,58^. All tracks were aligned to the hg38 reference genome.

### Statistical analysis

All statistical analyses were performed using GraphPad Prism and are reported in figure legends. *P*<0.05 was considered statistically significant.

## Data availability

BPNet models of human retinal cell types are available at https://zenodo.org/records/6330053. Model interpretation and TF-MoDISco outputs are available at https://zenodo.org/records/14061304. All other data supporting the findings of this study are available from the corresponding authors on reasonable request.

## ACKNOWLEDGEMENTS

This work was supported by the Knights Templar Eye Foundation (to S.K.W.), the Howard Hughes Medical Institute (to H.Y.C.), NIH R01-EY032585 (to S.W.), and NIH R21-EY035465 (to S.W.). Computing for this project was performed on the Sherlock cluster, a resource provided and maintained by the Stanford Research Computing Center.

## AUTHOR CONTRIBUTIONS STATEMENT

S.K.W., S.N., H.Y.C., and S.W. conceived the project. S.K.W., B.D., H.Y.C., and S.W. designed experiments. S.K.W., B.D., X.R., J.L., J.T., P.P., Z.L., C.N., Y.Z., S.H.S., A.D., R.M., Y.Q., Y.Z., J.Z., and S.W. performed experiments. S.K.W., B.D., S.N., S.H.K., and S.W. performed data analysis. S.K.W., Y.X., J.L.G., W.W., A.K., H.Y.C., and S.W. supervised the work. S.K.W., B.D., H.Y.C., and S.W. wrote the manuscript with input from all authors.

## REFERENCES

1. Wang, D., Tai, P. W. L. & Gao, G. Adeno-associated virus vector as a platform for gene therapy delivery. Nat. Rev. Drug Discov. 18, 358–378 (2019).

2. Russell, S. et al. Efficacy and safety of voretigene neparvovec (AAV2-hRPE65v2) in patients with RPE65-mediated inherited retinal dystrophy: a randomised, controlled, open-label, phase 3 trial. Lancet 390, 849–860 (2017).

3. Borel, F., Kay, M. A. & Mueller, C. Recombinant AAV as a platform for translating the therapeutic potential of RNA interference. Mol. Ther. 22, 692–701 (2014).

4. Ran, F. A. et al. In vivo genome editing using Staphylococcus aureus Cas9. Nature 520, 186–91 (2015).

5. Hudry, E. & Vandenberghe, L. H. Therapeutic AAV Gene Transfer to the Nervous System: A Clinical Reality. Neuron 101, 839–862 (2019).

6. Platt, R. J. et al. CRISPR-Cas9 knockin mice for genome editing and cancer modeling. Cell 159, 440–55 (2014).

7. Betley, J. N. & Sternson, S. M. Adeno-associated viral vectors for mapping, monitoring, and manipulating neural circuits. Hum. Gene Ther. 22, 669–77 (2011).

8. Hinderer, C. et al. Severe Toxicity in Nonhuman Primates and Piglets Following High-Dose Intravenous Administration of an Adeno-Associated Virus Vector Expressing Human SMN. Hum. Gene Ther. 29, 285–298 (2018).

9. Xiao, Y. et al. Circumventing cellular immunity by miR142-mediated regulation sufficiently supports rAAV-delivered OVA expression without activating humoral immunity. JCI insight 5, (2019).

10. Xiong, W. et al. AAV cis-regulatory sequences are correlated with ocular toxicity. Proc. Natl. Acad. Sci. U. S. A. 116, 5785–5794 (2019).

11. Clemens, P. R. et al. In vivo muscle gene transfer of full-length dystrophin with an adenoviral vector that lacks all viral genes. Gene Ther. 3, 965–72 (1996).

12. Brenner, M., Kisseberth, W. C., Su, Y., Besnard, F. & Messing, A. GFAP promoter directs astrocyte-specific expression in transgenic mice. J. Neurosci. 14, 1030–7 (1994).

13. Ye, G. J. et al. Cone-Specific Promoters for Gene Therapy of Achromatopsia and Other Retinal Diseases. Hum. Gene Ther. 27, 72–82 (2016).

14. Jüttner, J. et al. Targeting neuronal and glial cell types with synthetic promoter AAVs in mice, non-human primates and humans. Nat. Neurosci. 22, 1345–1356 (2019).

15. Mich, J. K. et al. Functional enhancer elements drive subclass-selective expression from mouse to primate neocortex. Cell Rep. 34, 108754 (2021).

16. Graybuck, L. T. et al. Enhancer viruses for combinatorial cell-subclass-specific labeling. Neuron 109, 1449–1464.e13 (2021).

17. Avsec, Ž., et al. Effective gene expression prediction from sequence by integrating long-range interactions. Nat. Methods 18, 1196–1203 (2021).

18. Avsec, Ž., et al. Base-resolution models of transcription-factor binding reveal soft motif syntax. Nat. Genet. 53, 354–366 (2021).

19. Linder, J., Srivastava, D., Yuan, H., Agarwal, V. & Kelley, D. R. Predicting RNA-seq coverage from DNA sequence as a unifying model of gene regulation. Nat. Genet. 57, 949–961 (2025).

20. Gosai, S. J. et al. Machine-guided design of cell-type-targeting cis-regulatory elements. Nature 634, 1211–1220 (2024).

21. DaSilva, L. F. et al. Designing synthetic regulatory elements using the generative AI framework DNA-Diffusion. Nat. Genet. (2025) doi:10.1038/s41588-025-02441-6.

22. Yin, C. et al. Iterative deep learning design of human enhancers exploits condensed sequence grammar to achieve cell-type specificity. Cell Syst. 16, 101302 (2025).

23. Watson, J. L. et al. De novo design of protein structure and function with RFdiffusion. Nature 620, 1089–1100 (2023).

24. Merchant, A. T., King, S. H., Nguyen, E. & Hie, B. L. Semantic design of functional de novo genes from a genomic language model. Nature (2025) doi:10.1038/s41586-025-09749-7.

25. Wang, S. K. et al. Single-cell multiome of the human retina and deep learning nominate causal variants in complex eye diseases. Cell Genomics 2, 100164 (2022).

26. Taskiran, I. I. et al. Cell-type-directed design of synthetic enhancers. Nature 626, 212–220 (2024).

27. de Almeida, B. P. et al. Targeted design of synthetic enhancers for selected tissues in the Drosophila embryo. Nature 626, 207–211 (2024).

28. Poché, R. A. et al. Lim1 is essential for the correct laminar positioning of retinal horizontal cells. J. Neurosci. 27, 14099–107 (2007).

29. Lin, C.-H. et al. Identification of cis-regulatory modules for adeno-associated virus-based cell-type-specific targeting in the retina and brain. J. Biol. Chem. 298, 101674 (2022).

30. Dalkara, D. et al. In vivo-directed evolution of a new adeno-associated virus for therapeutic outer retinal gene delivery from the vitreous. Sci. Transl. Med. 5, 189ra76 (2013).

31. Vrabec, J. P. & Levin, L. A. The neurobiology of cell death in glaucoma. Eye (Lond). 21 Suppl 1, S11–4 (2007).

32. Thoreson, W. B. & Mangel, S. C. Lateral interactions in the outer retina. Prog. Retin. Eye Res. 31, 407–41 (2012).

33. Macosko, E. Z. et al. Highly Parallel Genome-wide Expression Profiling of Individual Cells Using Nanoliter Droplets. Cell 161, 1202–1214 (2015).

34. Whipple, A. J. et al. Imprinted Maternally Expressed microRNAs Antagonize Paternally Driven Gene Programs in Neurons. Mol. Cell 78, 85–95.e8 (2020).

35. Zhang, Y. et al. Fast and sensitive GCaMP calcium indicators for imaging neural populations. Nature 615, 884–891 (2023).

36. Goetz, J. et al. Unified classification of mouse retinal ganglion cells using function, morphology, and gene expression. Cell Rep. 40, 111040 (2022).

37. Saito, M. et al. Diphtheria toxin receptor-mediated conditional and targeted cell ablation in transgenic mice. Nat. Biotechnol. 19, 746–50 (2001).

38. Sluch, V. M. et al. Differentiation of human ESCs to retinal ganglion cells using a CRISPR engineered reporter cell line. Sci. Rep. 5, 16595 (2015).

39. Dyer, M. A., Livesey, F. J., Cepko, C. L. & Oliver, G. Prox1 function controls progenitor cell proliferation and horizontal cell genesis in the mammalian retina. Nat. Genet. 34, 53–8 (2003).

40. Dorgau, B. et al. Deciphering the spatiotemporal transcriptional and chromatin accessibility of human retinal organoid development at the single-cell level. iScience 27, 109397 (2024).

41. Chan, K. Y. et al. Engineered AAVs for efficient noninvasive gene delivery to the central and peripheral nervous systems. Nat. Neurosci. 20, 1172–1179 (2017).

42. Tabebordbar, M. et al. Directed evolution of a family of AAV capsid variants enabling potent muscle-directed gene delivery across species. Cell 184, 4919–4938.e22 (2021).

43. Coughlin, G. M. et al. Spatial genomics of AAV vectors reveals mechanism of transcriptional crosstalk that enables targeted delivery of large genetic cargo. Nat. Biotechnol. (2025) doi:10.1038/s41587-025-02565-4.

44. Agarwal, V. et al. Massively parallel characterization of transcriptional regulatory elements. Nature 639, 411–420 (2025).

45. Zhang, K. et al. A single-cell atlas of chromatin accessibility in the human genome. Cell 184, 5985–6001.e19 (2021).

46. Liu, B. B. et al. Dissecting regulatory syntax in human development with scalable multiomics and deep learning. (2025) doi:10.1101/2025.04.30.651381.

47. Feng, J., Liu, T., Qin, B., Zhang, Y. & Liu, X. S. Identifying ChIP-seq enrichment using MACS. Nat. Protoc. 7, 1728–40 (2012).

48. Tan, G. & Lenhard, B. TFBSTools: an R/bioconductor package for transcription factor binding site analysis. Bioinformatics 32, 1555–6 (2016).

49. Shrikumar, A. et al. Technical Note on Transcription Factor Motif Discovery from Importance Scores (TF-MoDISco) version 0.5.6.5. (2018).

50. Bruse, N. & Heeringen, S. J. van. GimmeMotifs: an analysis framework for transcription factor motif analysis. bioRxiv 474403 (2018) doi:10.1101/474403.

51. Azim, E., Jiang, J., Alstermark, B. & Jessell, T. M. Skilled reaching relies on a V2a propriospinal internal copy circuit. Nature 508, 357–63 (2014).

52. Wu, J. et al. Transcardiac Perfusion of the Mouse for Brain Tissue Dissection and Fixation. Bio-protocol 11, e3988 (2021).

53. Stuart, T. et al. Comprehensive Integration of Single-Cell Data. Cell 177, 1888–1902.e21 (2019).

54. Griffiths-Jones, S., Grocock, R. J., van Dongen, S., Bateman, A. & Enright, A. J. miRBase: microRNA sequences, targets and gene nomenclature. Nucleic Acids Res. 34, D140–4 (2006).

55. Brainard, D. H. The Psychophysics Toolbox. Spat. Vis. 10, 433–6 (1997).

56. Acarón Ledesma, H., et al. Dendritic mGluR2 and perisomatic Kv3 signaling regulate dendritic computation of mouse starburst amacrine cells. Nat. Commun. 15, 1819 (2024).

57. Luo, Z. et al. An Optimized System for Effective Derivation of Three-Dimensional Retinal Tissue via Wnt Signaling Regulation. Stem Cells 36, 1709–1722 (2018).

58. Li, D., Hsu, S., Purushotham, D., Sears, R. L. & Wang, T. WashU Epigenome Browser update 2019. Nucleic Acids Res. 47, W158–W165 (2019).

