## Supplementary Information for "Deep learning-guided design of cell type-specific AAV promoters"

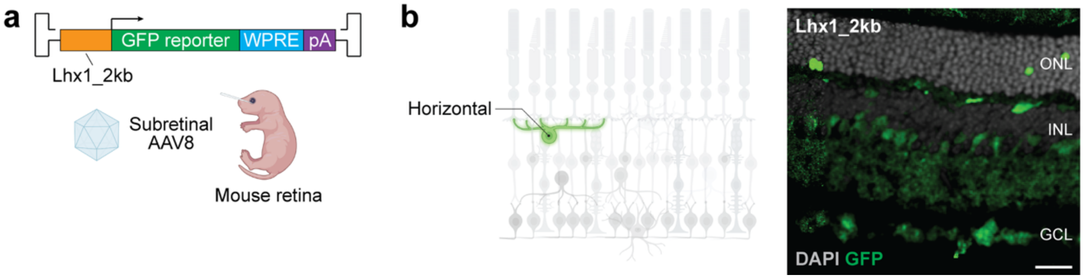


**Supplementary Figure 1. Activity of a native horizontal cell promoter sequence.**

**a,** Schematic of native promoter experiments. The 2 kb sequence immediately upstream of the mouse *Lhx1* 5' untranslated region was placed upstream of a GFP reporter sequence, packaged in the AAV8 capsid, and subretinally injected into neonatal mice to target horizontal cells (HCs) in the retina. **b,** *Left:* Schematic of HCs. *Right:* Lhx1_2kb promoter activity in mouse retina three weeks after AAV8 transduction. n = 8 eyes. Scale bar, 20 µm. ONL, outer nuclear layer; INL, inner nuclear layer; GCL, ganglion cell layer.

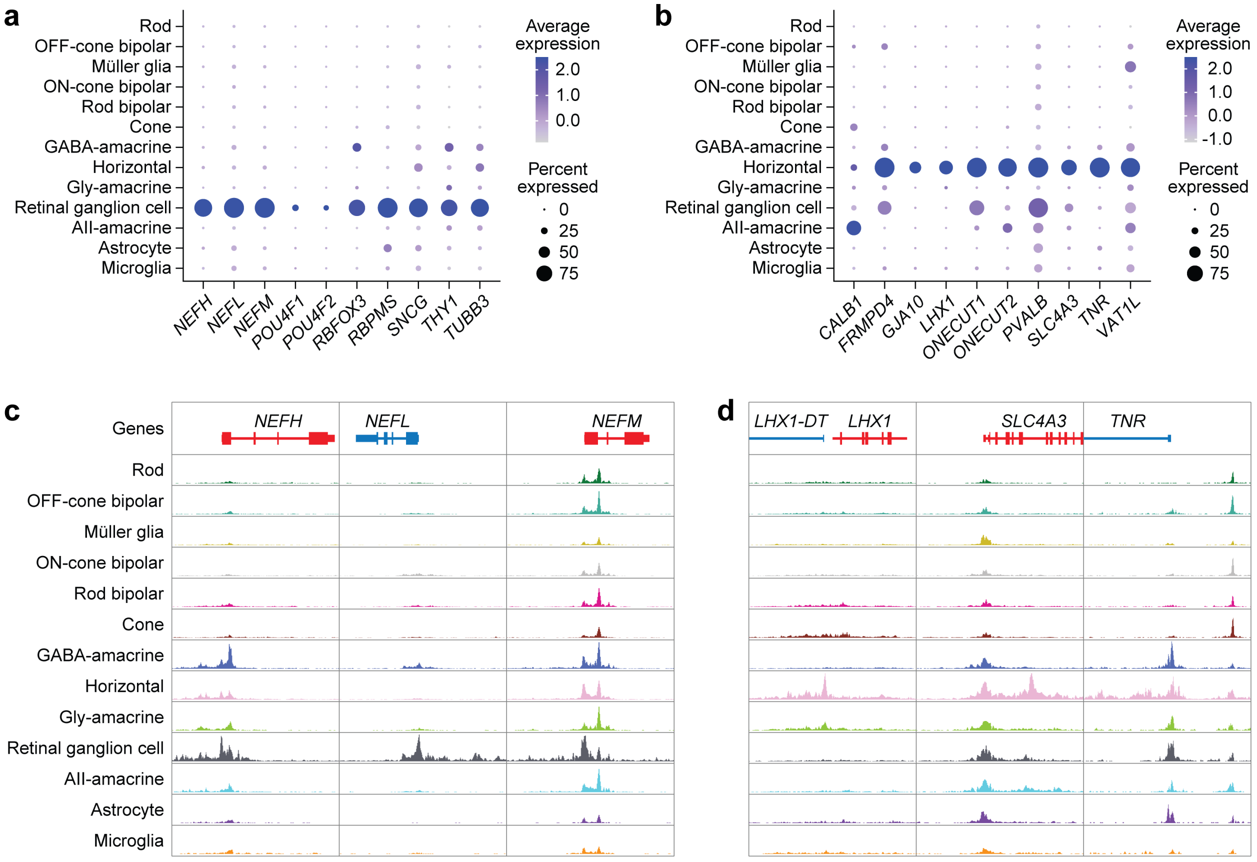


**Supplementary Figure 2. Comparison of cell type marker genes.**

**a-b,** Dot plot visualizing the normalized RNA expression of putative retinal ganglion cell (RGC) (a) or HC (b) marker genes in human retinal cell types[25]. The color and size of each dot correspond to the average expression level and fraction of expressing cells, respectively. **c-d,** Sequencing tracks of cell type chromatin accessibility near selected RGC (c) or HC (d) marker genes. Each track depicts the aggregate scATAC-seq signal in the given cell type normalized by the number of reads in transcriptional start site (TSS) regions. Coordinates for each region: *NEFH* (chr22:29475296-29491829), *NEFL* (chr8:24949448-24964533), *NEFM* (chr8:24907410-24921098), *LHX1* (chr17:36929441-36945474), *SLC4A3* (chr2:219622838-219634611), *TNR* (chr1:175733110-175753171).

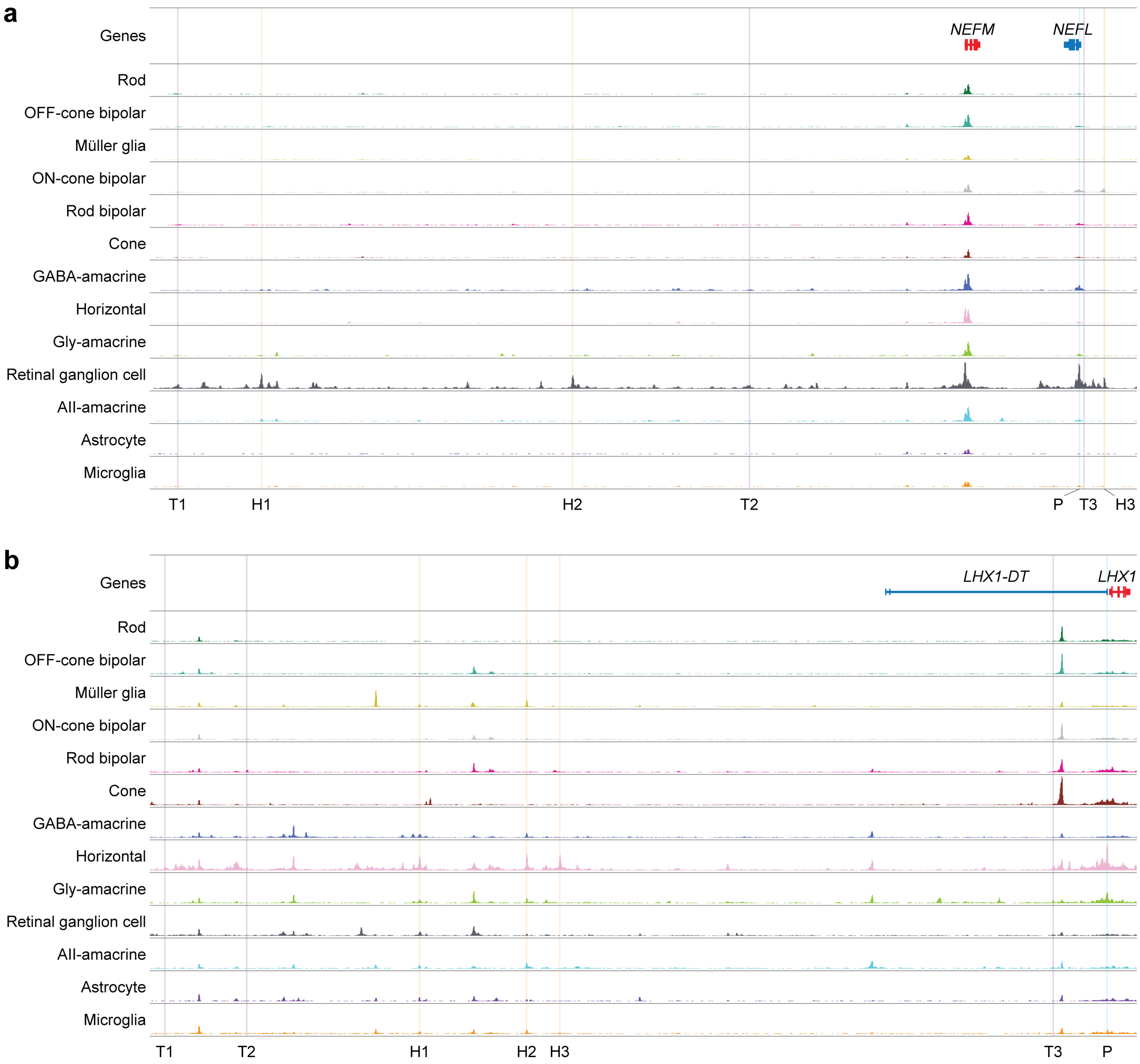


**Supplementary Figure 3. Endogenous candidate enhancers targeting retinal ganglion cells and horizontal cells.**

**a-b,** Sequencing tracks of chromatin accessibility in human retinal cell types. Vertical lines indicate locations of the base promoter (P), highly accessible regions (H1-H3), and transcription factor binding site (TFBS)-enriched peaks (T1-T3) selected to target RGCs (a) or HCs (b). Coordinates for each region: chr8:24607917-24978286 (a), chr17:36612867-36946800 (b).

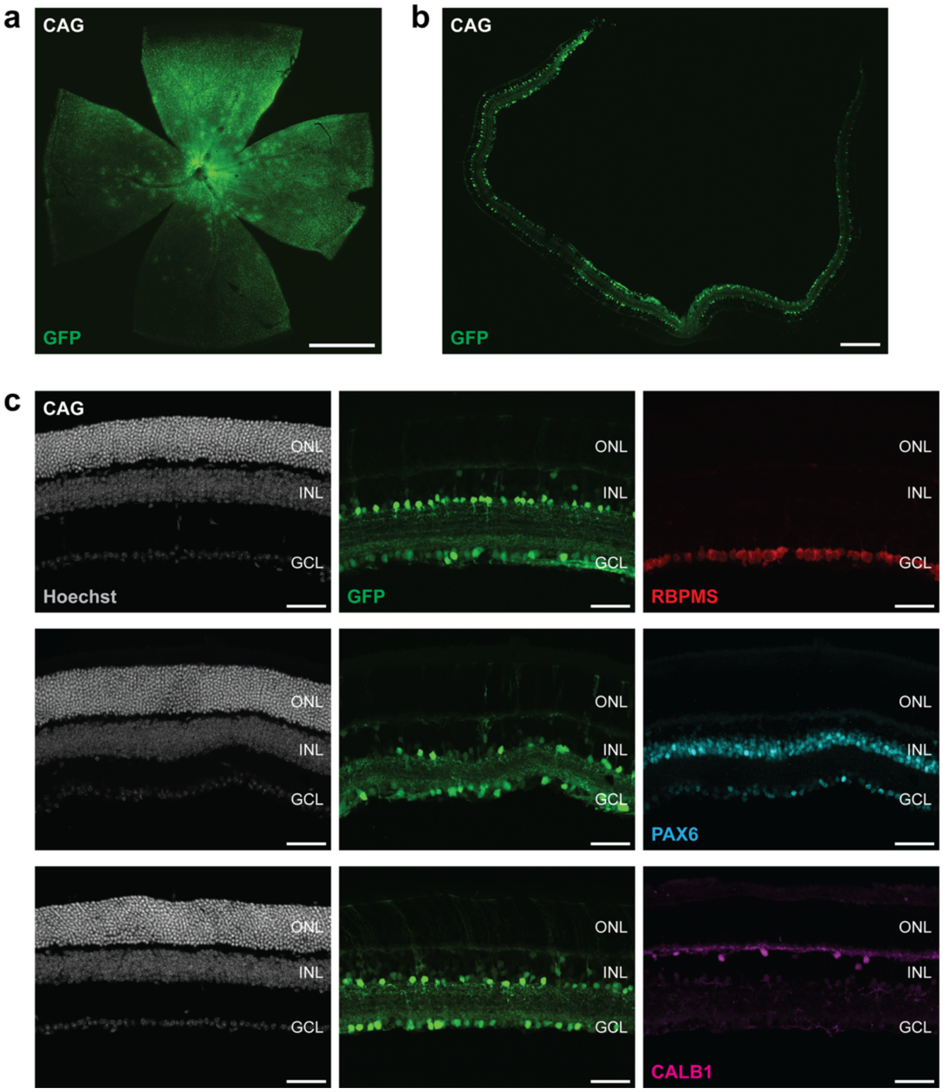


**Supplementary Figure 4. AAV7m8 transduction of retinal cell types.**

**a-b,** Mouse retinal flat-mount (a) and section (b) three weeks after intravitreal injection of an AAV7m8 vector expressing GFP under the control of the CAG promoter. n = 3 eyes. Scale bars, 1 mm (a), 250 µm (b). **c,** CAG promoter activity and immunostaining for cell type markers in mouse retina three weeks after AAV7m8 transduction. n = 3 eyes. Scale bars, 50 µm. ONL, outer nuclear layer; INL, inner nuclear layer; GCL, ganglion cell layer.

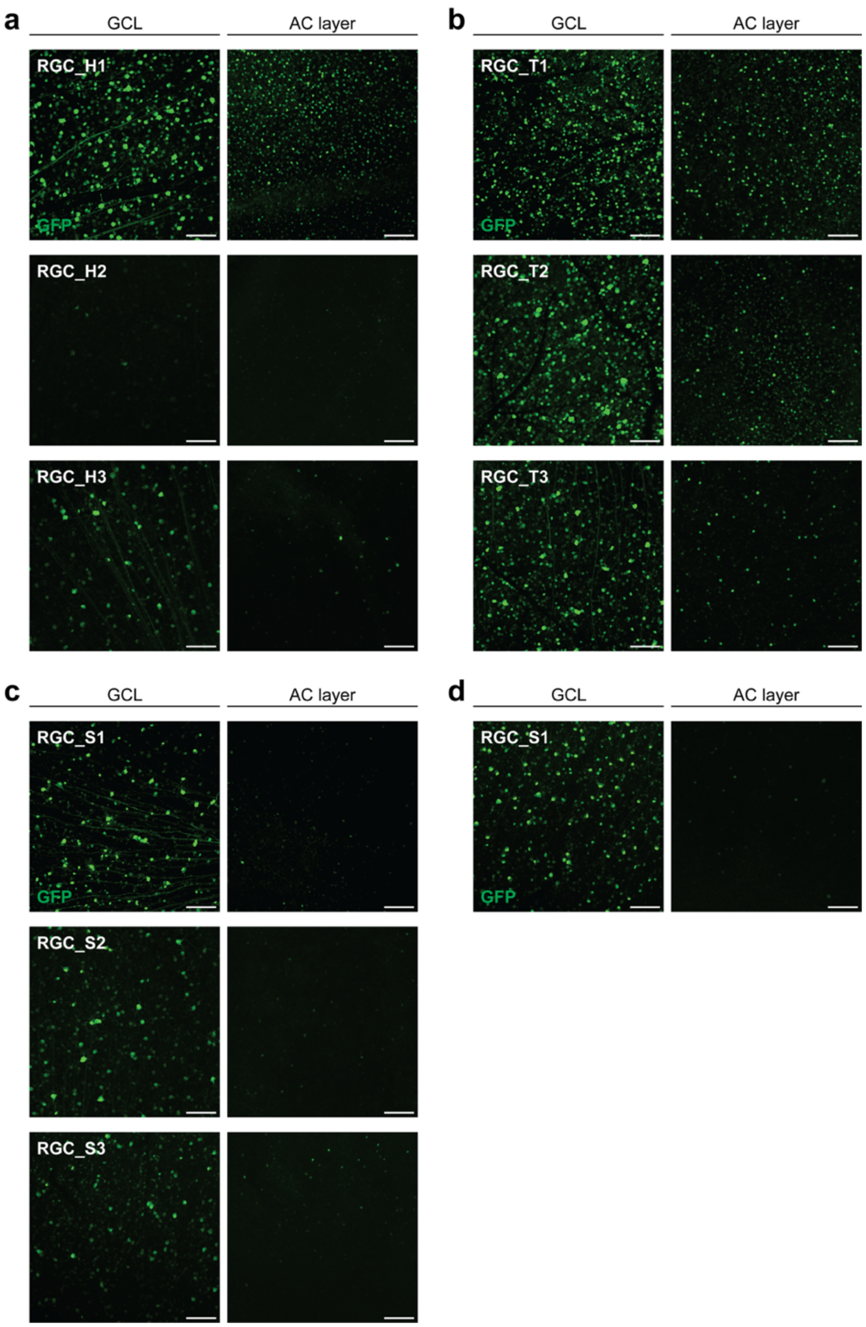


**Supplementary Figure 5. Activity of AAV promoters targeting retinal ganglion cells.**

**a-d,** Activity of the indicated AAV promoters in the ganglion cell layer (GCL) and amacrine cell (AC) layer of flat-mounted mouse retina three weeks after AAV7m8 (a-c) or AAV2 (d) transduction. n = 3-6 eyes per group. Scale bars, 100 µm.

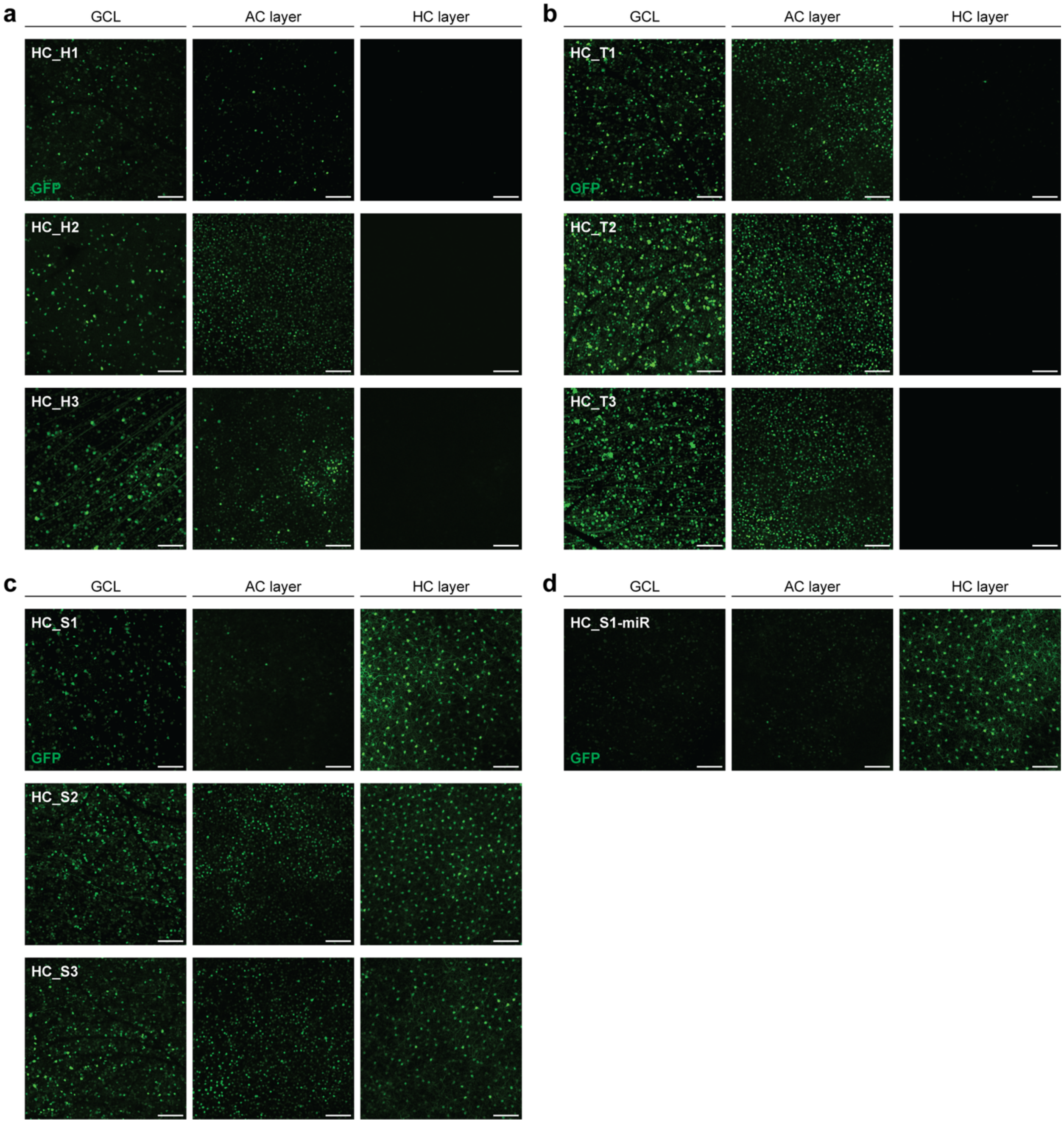


**Supplementary Figure 6. Activity of AAV promoters targeting horizontal cells.**

**a-d,** Activity of the indicated AAV promoters in the GCL, AC layer, and HC layer of flat-mounted mouse retina three weeks after AAV7m8 transduction. n = 3-6 eyes per group. Scale bars, 100 μm.
